# Predictive processing in biological motion perception: Evidence from human behavior

**DOI:** 10.1101/2024.02.03.578729

**Authors:** Hüseyin O. Elmas, Sena Er, Ada D. Rezaki, Aysesu Izgi, Buse M. Urgen, Huseyin Boyaci, Burcu A. Urgen

## Abstract

Biological motion perception plays a crucial role in understanding the actions of other animals, facilitating effective social interactions. While foundation of biological motion perception is rooted in bottom-up processes, as evidenced by point-light display studies, real-world complexities necessitate the involvement of top-down processes, such as attention and expectation. This study investigates the impact of expectations on biological motion perception using a cued individuation task with point-light display stimuli. We conducted three experiments, each providing advance information about distinct aspects of the subsequent biological motion stimuli – specifically information about action, emotion, and gender. Our results revealed a pattern in the action experiment, where participants demonstrated significantly slower response times for incongruent trials than congruent ones, but only under the 75% cue validity condition. This effect was notably absent in the emotion and gender experiments. Our exploration underscores the multi-faceted nature of biological motion perception, highlighting that while the brain adeptly harnesses prior cues to anticipate and interpret stimuli, the nature and reliability of these cues play a pivotal role on their effects. Specifically, action-related information stands out as an important modulator, possibly due to its evolutionary significance and unique neural processing pathway. These findings not only agree with the principles of predictive processing but also pave the way for future research, emphasizing the need to utilize naturalistic, complex stimuli together with neuroimaging methods to create more comprehensive models of biological motion perception.

## 1. Introduction

The ability to perceive and understand the actions of other animals is pivotal in facilitating accurate and reliable social interactions (Blake and Shiffrar, 2007). The evolutionary and social significance of biological motion perception has led animals to develop advanced processing mechanisms. These mechanisms enable swift and robust inferences from cues in biological motion (Simion et al., 2008; De Agrò et al., 2021; Troje and Basbaum, 2008; Omori and Watanabe, 1996), suggesting an innate ability for biological motion perception. Research utilizing point-light display (PLD) stimuli supports the automatic, bottom-up, innate nature of biological motion perception (Johansson, 1973; Troje and Basbaum, 2008). Additionally, roots of biological motion perception can be traced back to the early developmental stages of animals. For instance, two-day-old humans already show a preference for PLD’s of biological motion over inanimate objects (Simion et al., 2008), and by age five, they achieve discrimination capabilities that match levels of adults (Blake and Shiffrar, 2007). This innate inclination isn’t exclusive to humans; even newborn chicks, without any prior visual exposure, are drawn to biological motion (Vallortigara et al., 2005).

Further emphasizing the foundational nature of biological motion perception, Mather et al. (1992) found that perception of the PLD walker was dependent on low-level visual mechanisms that operate over small spatial and temporal windows by manipulating the time intervals between each frame of a PLD walker. Reinforcing this perspective, models like those proposed by Giese and Poggio (2003), which focus exclusively on this bottom-up approach, align remarkably well with human performance across various scenarios. While the foundational role of bottom-up processes in biological motion perception is clear, the complexity of real-world scenarios necessitates the involvement of top-down processes on various kinds of perception (Gilbert and Li, 2013; De Lange et al., 2018; Spaak et al., 2022; Peelen et al., 2023). In situations with ambiguous or degraded stimuli, mere bottom-up cues might be insufficient and top-down processes like attention and expectation also affect perception (Blake and Shiffrar, 2007). Supporting this perspective, several studies underscored the importance of attention in biological motion perception. Cavanagh et al. (2001) demonstrated that distinguishing biological motion amidst other distractors necessitates attention. Similarly, Thornton et al. (2002) emphasized the role of attention in detecting PLDs when bottom-up processing is compromised, yet found no significant effect of attention when bottom-up cues were clear. Parasuraman et al. (2009) observed that sustained attention to a demanding task could diminish the ability to detect biologival motion targets, but this effect was pronounced only in visually degraded conditions. On a neural level, Hirai et al. (2005) explored the effects of attention on biological motion sensitive ERP components, identifying the N330, a negative ERP component emerging approximately 330ms post-stimulus, as being modulated by attention. This finding further solidifies the argument that biological motion perception is influenced by both bottom-up and top-down processes.

Just as attention profoundly influences our perception, expectation plays an equally important role (Gilbert and Li, 2013; Aitken et al., 2020). The impact of expectation on perception is multifaceted, becoming especially pronounced when faced with weak or ambiguous sensory input. For example, when expecting a particular stimulus, the decision-making criteria in tasks may shift (Bang and Rahnev, 2017). Similarly, such expectations can refine how the brain represents the anticipated stimulus (Kok et al., 2012; Cheadle et al., 2015; Kok et al., 2017).

Most research on the effects of visual expectation has primarily employed low-level stimuli like gabor filters, randomly moving dots, or simple lines. These stimuli have been pivotal in isolating the effects of expectation and offering insights into predictive processing mechanisms. However, their ecological validity beyond laboratory settings is often questioned, as it’s unclear if the same expectation effects are observable with more complex, socially relevant stimuli. Notably, studies using faces and houses to investigate the effect of expectation on perceptual-decision making, such as those by Urgen and Boyaci (2021), indicate that violated expectations can slow down sensory processing. Similarly, research by De Loof et al. (2016) demonstrates that incongruent predictive information can slow down decision-making by altering decision thresholds. Yet, there remains a gap in understanding how these mechanisms translate to the perception of biological motion, which is crucial given the dynamic nature of our environment and the movements of other agents. Our study aims to address this gap by building on the findings of De Loof et al. (2016) and Urgen and Boyaci (2021). We seek to apply a similar paradigm to biological motion perception and extend these insights to explore how expectation mechanisms function within the context of biological motion perception. Thus, there is a clear imperative to explore the impact of expectation on biological motion perception to further unravel the complexities of this cognitive process.

Existing research on biological motion and expectation suggests an influence of expectation on biological motion perception. For instance, Hunt and Halper (2008) discovered that when point lights are replaced with objects, the stimulus becomes ambiguous. Naive observers struggle to perceive biological motion even when the motion and form information remain intact. However, observers who were informed about the presence of biological motion in the stimuli could reliably identify it. Furthermore, even when motion information is manipulated to a level that only bottom-up processing should not yield a meaningful interpretation, observers can still perceive biological motion (Thornton et al., 1998; Bülthoff et al., 1998). These findings suggest that in certain circumstances where bottom-up information is ambiguous, top-down processes like expectation and attention influence biological motion processing. Template-matching models of biological motion perception, which incorporate top-down processes based on form information, have been proposed and successfully explain biological motion recognition performance (Lange et al., 2006). Action perception studies employing naturalistic action videos and expectation paradigms have provided insights into the underlying mechanisms, as evidenced by the findings of (Saygin et al., 2012; Urgen et al., 2018; Urgen and Saygin, 2020). However, there remains an absence of studies that behaviorally interrogate biological motion perception, particularly in the context of probabilistic cuing paradigms observed in aforementioned research using relatively simpler stimulus. The imperative for such an exploration has been previously suggested by (Urgen and Miller, 2015). Therefore, there’s a gap in current research that explores the impact of expectation on biological motion perception while also using a task independent of the expectation.

Furthermore, biological motion is a multifaceted stimulus, offering a wealth of information that can serve as prior knowledge. These diverse sources of prior information might have varied impacts on biological motion perception. Different types of prior knowledge, such as the type of action performed by the agent, the gender or the emotional state of the moving agent, might be processed by distinct neural populations. For example, research suggests that the posterior parietal cortex processes information related to observed action of the the agent, anterior temporal pole and fusiform areas processes gender of the agent, and amygdala processes emotional state of the agent (Orban, 2018; Phelps, 2006; Kaul et al., 2011). Further research in this area is essential to refine our understanding and develop comprehensive models of biological motion perception that also accounts for different types of information.

To address the existing gap in the literature, we explored the influence of prior information on biological motion perception using a cued individuation task, building upon methodologies from earlier studies (De Loof et al., 2016; Urgen and Boyaci, 2021). In our experiments, participants were presented with a cue, offering predictive information about the subsequent biological motion stimulus. Following this cueing phase, participants were exposed to point-light display stimuli, encompassing both a biological motion and its scrambled counterpart. Participants were asked to indicate the location of the biological motion while we measured their reaction time and accuracy. This paradigm was repeated across three experiments, each varying in the type of prior information and biological motion stimuli presented. Within each experiment, both the congruency and validity of the cues were manipulated.

Drawing from existing research on predictive processing and biological motion perception, we hypothesize that biological motion stimuli to be individuated slower following incongruent cues compared to biological motion stimuli following congruent cues. Additionally, we anticipate that different categories of information, such as gender, emotion and action information, would influence biological motion perception in varied ways. More specifically, we predict that the most critical and relevant information would exert a stronger effect on biological motion perception. Furthermore, the reliability of the prior information is expected to shape its overall impact on biological motion perception.

## 2. Methods and Materials

### 2.1. Participants

60 participants were recruited for three experiments: For the action experiment, we had 21 participants (14 female, ages: 22.00*±*2.07). The emotion experiment included 19 participants (15 female, ages: 21.15 *±* 3.25). For the gender experiment, there were 21 participants (12 female, ages: 21.33*±*2.78). All participants either had normal vision or vision corrected to normal and had not been on any long-term neurological or psychiatric medications. The study received approval from Bilkent University’s Human Research Ethics Committee. All participants provided written informed consent and completed a pre-screening form before the experiments. In each experiment, participants took part in four sessions for different validity conditions (Neutral, 50%, 75% and 100%) on separate days, with the session order randomized. They were trained on the stimuli and task before the first session. After completing all sessions, participants were compensated with a monetary reward.

### 2.2. Stimuli and Apparatus

Our study used two types of visual stimuli: central cues during the cue period and peripheral point-light displays (PLDs) in the subsequent target period. The cues appeared centrally on the screen, while the PLDs, both intact and scrambled versions, were positioned on the screen’s left and right sides. These stimuli were presented in separate phases of a trial against a uniform gray background. To manipulate the type of prior information, we used three distinct cues (refer to Fig. 1) and PLDs across the experiments. The cue, sized at 3.5 x 3.5 visual degrees, was centrally displayed for 2 seconds. In contrast, the PLDs, measuring 5.2 x 8.3 visual degrees, were placed 7 visual degrees from the screen’s center.

**Figure 1:**
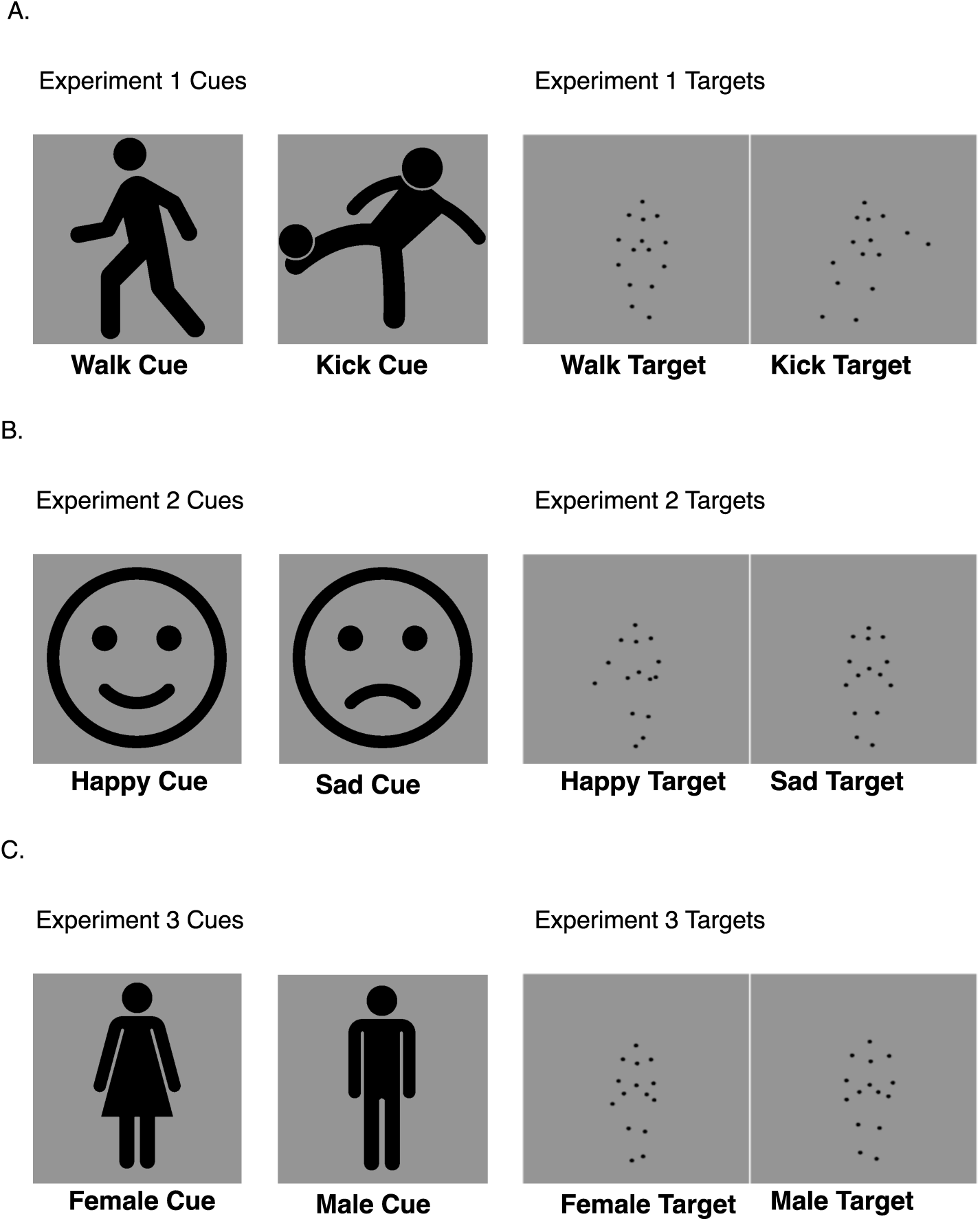
Cues and corresponding target biological motion stimuli used in the experiments. Cues are on the two left columns, and targets are on the right two columns. A. Experiment 1: Action cues and corresponding PLD targets, B. Experiment 2: Emotion cues and corresponding PLD targets, C. Experiment 3: Gender cues and corresponding PLD targets

For our PLD stimuli, we sourced initial selections from the Carnegie Mellon University’s motion capture database (CMU Graphics Lab, (Hodgins, 2015)) as shown in Figure 1, A. We then tailored these motion capture stimuli to our experimental design using the Motion Kinematic and Kinetic Analyzer (MOKKA, 2018, Barre and Armand (2014)). For the emotion-focused experiment, we chose sad and happy walking actions from the same database (Ma et al., 2006) (see Figure 1,B). For the gender experiment, we crafted new stimuli. We used the walking action from the action experiment as a foundational template to ensure consistency across experiments. This walking action was then adapted by adjusting specific point distances and positions to produce discernible male and female cues. Specifically we used the form and motion patterns from Troje and colleagues work and their accompaniying demo BML Walker (Troje, 2002, 2008).

To validate the efficacy of the PLD stimuli in conveying gender and emotion cues, a two-phase online survey was administered via Qualtrics (Qualtrics, 2023). In the initial phase, participants viewed videos of the stimuli set against a neutral gray background. Participants were then asked to rate the gender (ranging from Female to Male) or emotion (from Sad to Happy) of the stimuli on a 5-point Likert scale, spanning -2 to 2. Subsequently, in the second phase, participants were shown paired actions (happy/sad and male/female) and were instructed to identify the position (left or right) of a specified stimulus (e.g., “Identify the side displaying the male walker”). 113 completed the survey. During the first phase, the stimuli received the following mean scores on the -2 to 2 scale: Female (M=-0.699, SD=1.093), Male (M=0.858, SD=1.068), Sad (M=-1.265, SD=0.876), and Happy (M=- 0.54, SD=1.536). Dependent sample t-tests revealed significant disparities in scores between the male-female and happy-sad stimulus pairs (*p <* 0.001). In the second phase, for every query spanning the two aforementioned categories, the accuracy surpassed the chance level of 0.5 (all *p <* 0.001. The results demonstrated that the PLD stimuli effectively convey gender and emotion cues, with participants consistently identifying these cues at levels significantly above chance.

We generated a set of scrambled motion stimuli by randomly shuffling the starting positions of each of the dots in the original biological motion stimuli using the BioMotion Toolbox (van Boxtel and Lu, 2013). It’s crucial to note that while we changed the initial locations of these dots, their individual motion paths were left unchanged. This method disrupted the coherent global motion pattern seen in the intact biological motion stimuli. However, the number of points and local motion patterns remained consistent.

### 2.3. Experiment Design and Procedure

Our experimental methodology involved conducting a series of three distinct experiments, each tailored to manipulate the nature of the prior information provided to participants. This manipulation centered around three specific types of cues, in conjunction with point-light display (PLD) stimuli. The cues were designed to convey distinct categories of information: the action being performed (such as kicking or walking), the emotional state (either happy or sad), and the gender (female or male) of the figures represented in the PLD stimuli. Participants in each experiment undertook a cued individuation task, adapted for biological motion stimuli from (De Loof et al., 2016; Urgen and Boyaci, 2021). Trials started with a cue symbol presentation, predicting the upcoming biological motion stimulus. Following the cue, an intact (BM) and scrambled version (SM) of a point-light biological motion stimulus appeared on the screen’s left and right sides for 1.2 seconds (Figure 2). Participants’ task was to identify the biological motion stimulus’s location, responding with the appropriate arrow key. Participants could respond as soon as the PLD stimuli are presented and in a 2-seconds response period proceeding after the stimulus. Notably, while cues pertained to the BM stimulus’s features, the task was spatial, ensuring cue information and task objective were independent. Following their response, participants received feedback on their accuracy for 2 seconds. To maintain task difficulty and promote reliance on prior information, we incorporated noise dots with the biological motion (BM) and scrambled motion (SM) stimuli (Figure 2). Quest (Watson and Pelli, 1983), an adaptive psychometric procedure, ensured consistent task difficulty across participants and sessions by adjusting the number of noise dots depending on the performance of the participant. Both cues and biological motion stimuli were randomized and balanced for each participant, as was the screen position of the biological motion stimulus.

**Figure 2:**
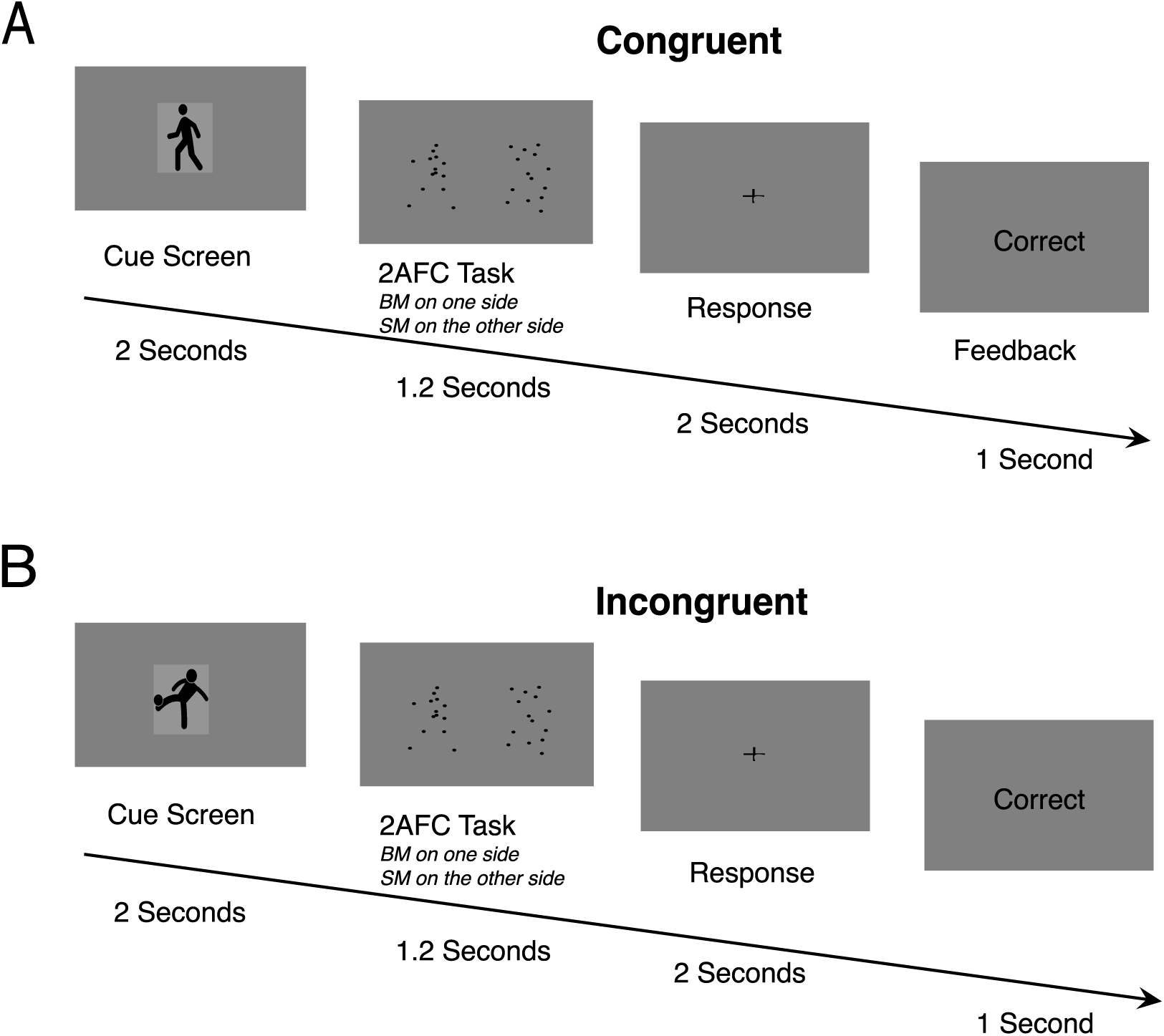
The trial structure and comparison between incongruent and congruent trials. A. A congruent trial B. An incongruent trial.

**Figure 3:**
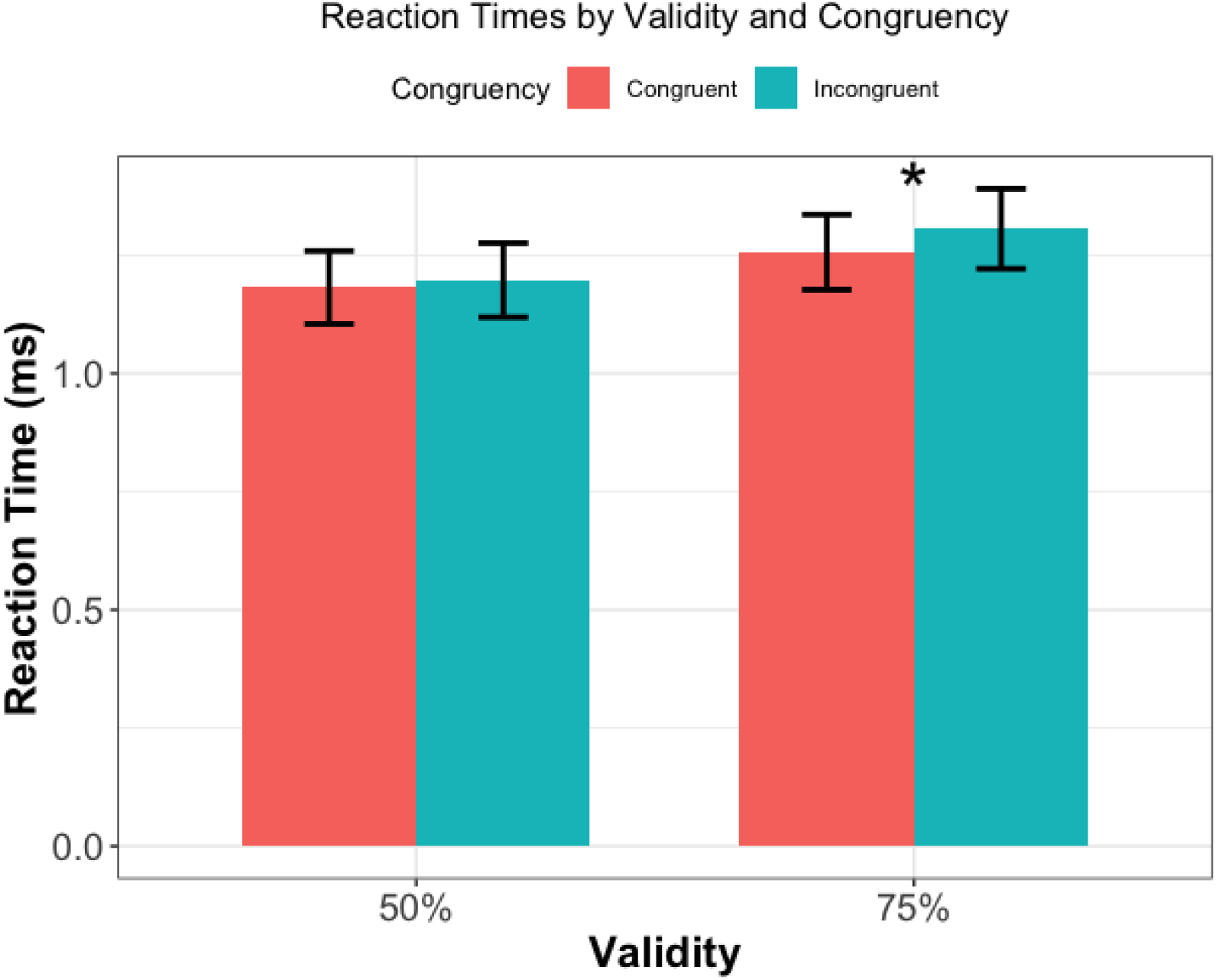
Reaction times for congruent and incongruent conditions when the validity is 50% and 75% in action experiment. * Denoting statistical significance (p=0.025)

**Figure 4:**
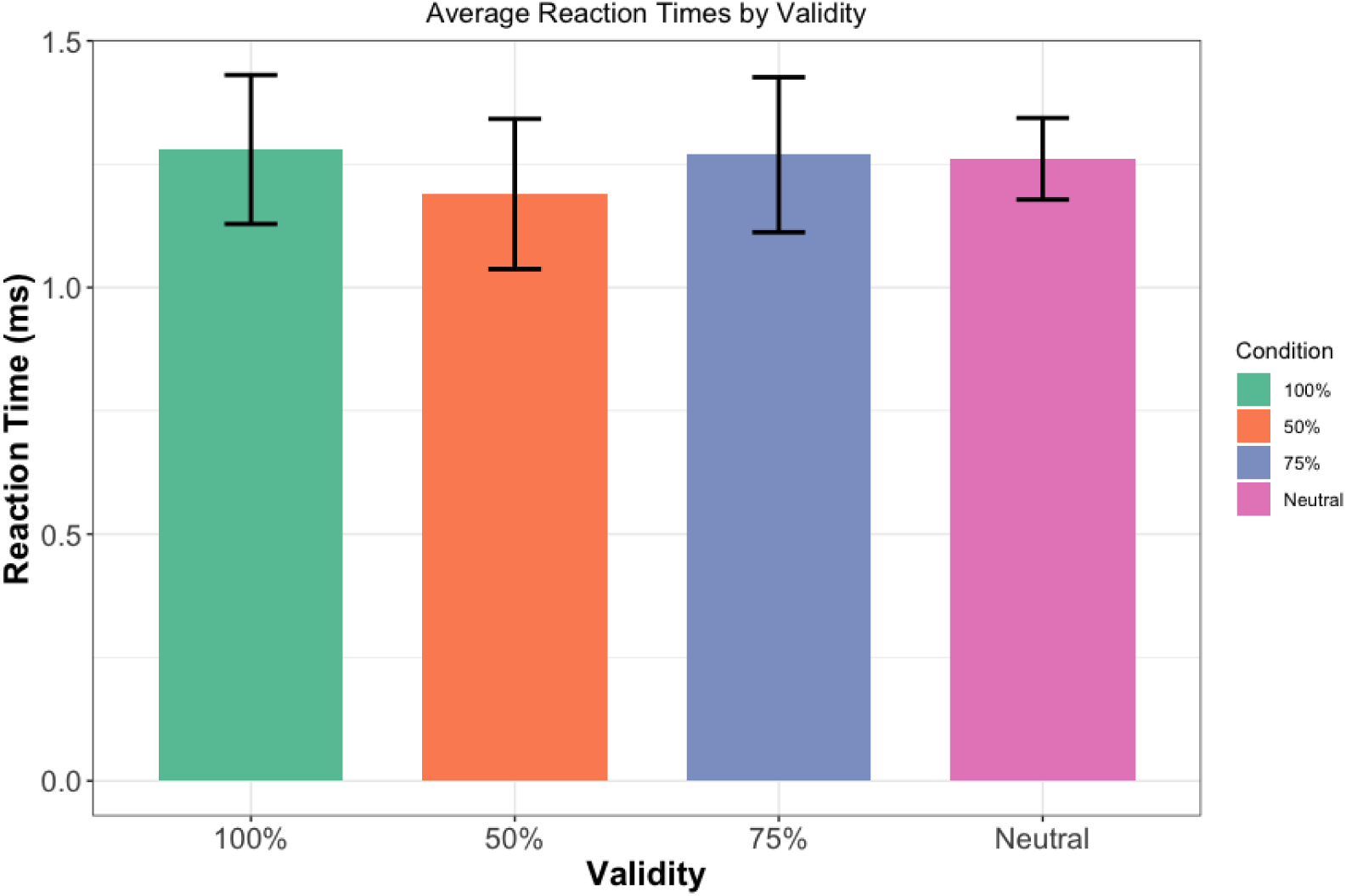
Overall reaction times for different validity conditions in action experiment

Within each experiment, we adjusted both the congruency and validity of the cues. Trials were classified as either congruent or incongruent based on the alignment between the cue and the subsequent stimulus. In congruent trials, the preceding cue matched the PLD stimuli’s features, whereas in incongruent trials, they differed. For instance, if a ”kick” symbol was the cue, congruent trials would display a ”kick” PLD alongside a scrambled PLD (see figure 2). Each experiment spanned four sessions. In three of these, the congruency ratio (or ”validity”) of the cues varied (50%, 75%, and 100%). The fourth was a neutral session, using a question mark as the cue. In addition to this neutral condition, we included a 50% validity condition as a control condition. In this scenario, the cue presented to participants holds no informative value over time, effectively serving as a non-informative cue with a chance alignment of 50%. This allows for two control condition a completely neutral cue and one that, while present, does not provide reliable predictive information consistently.

Each session was divided into three blocks, totaling 288 trials. Before starting the first session, participants were introduced to the biological motion stimuli and briefed on the task. Participants who couldn’t achieve a 70% accuracy rate in a noise-free training session were excluded from the study. Ahead of subsequent sessions, participants received a refresher on the task and stimuli, viewing the biological motion stimuli and completing noisy practice trials. They were also briefed on the cue validity for the upcoming session. During these training phases, cues were consistently informative and congruent.

Sessions took place in the psychophysics laboratory at Bilkent University, Aysel Sabuncu Brain Research Center. Participants were seated 57 cm from a CRT screen (HP P1230, 22 inches, 85 Hz). Participants were instructed to maintain focus on a central fixation cross throughout the task. To ensure consistent viewing distance and comfort, an adjustable chin-rest supported participants’ heads.

### 2.4. Data Analysis

In our analysis, we aimed to understand how reaction times changes when the presented biological motion stimulus aligns (congruent) or conflicts (incongruent) with the preceding cue. To this end, for the 50% and 75% cue validity conditions, we conducted a 2 (Validity) x 2 (Congruency) repeated-measures ANOVA, independently for each cue type. Note that, validity fac-tor includes only 50% AND 75% conditions since congruency is only valid for those two conditions. Additionally, to evaluate the effect of cue validity, we also carried out a one-way ANOVA on mean reaction times across cue validities (Neutral, 50%, 75%, and 100%), independently for each cue type.

## 3. Results

### 3.1. Action Experiment

In the action experiment, our analysis revealed a main effect of congruency on reaction times (F(1,20)=5.828, p=0.025, *η*^2^ = 0.226). Specifically, we found that participants responded significantly faster in congruent trials than in incongruent ones. However, there was no main effect of validity (F(1,20)=1.56, p=0.2). Finally, there was a marginally significant interaction effect between validity and congruency (F(1,20)=3.720, p=0.068) (Figure: 3). Post-hoc analysis of the aforementioned ANOVA revealed that the effects of congruency were significant only within the 75% validity condition.

We found the average reaction times across different validity conditions were as follows: 1.261 (SD=0.193) for neutral, 1.190 (SD=0.356) for 50% validity, 1.269 (SD=0.367) for 75% validity, and 1.280 (SD=0.353) for 100% validity. A one-way ANOVA comparing reaction times across these validity levels revealed no main effect of validity on reaction times(F(3,60)=0.796, p=0.5). (Figure:4).

To recapitulate the results of our experiment with action cues, participants exhibited slower response times when presented with incongruent cues compared to congruent cues in the 75% validity session. Notably, this difference was observed exclusively in the 75% validity session, wherein the cue provided reasonably reliable information. In contrast, no such difference was noted in the 50% validity session, where the cues did not offer any prior information about the action of the upcoming biological motion stimulus.

### 3.2. Emotion Experiment

For the emotion experiment, our 2 (Congruency) x 2 (Validity) ANOVA revealed no main effect for congruency (F(1,18) = 0.124, p = .728) or for validity (F(1,18) = 0.124, p = .728). Additionally, the interaction effect between congruency and validity was not significant (F(1,18) = 0.544, p = 0.470), see figure 5.

**Figure 5:**
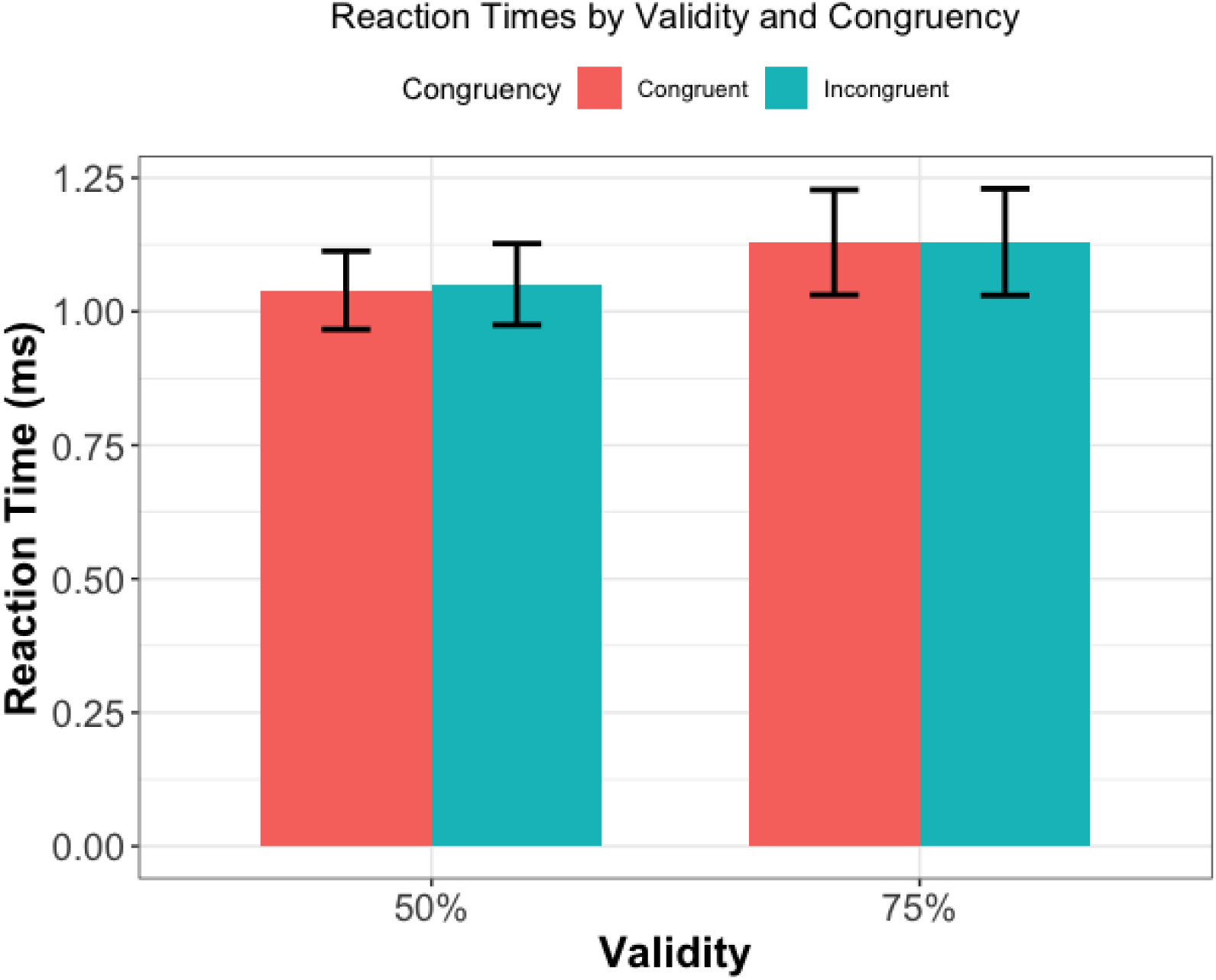
Reaction times for congruent and incongruent conditions when the validity is 50% and 75% in emotion experiment

A one way ANOVA across validity levels in the emotion experiment, showed no effect of validity on reaction times (F(3,60)=0.796, p=0.5) (Figure 6).

**Figure 6:**
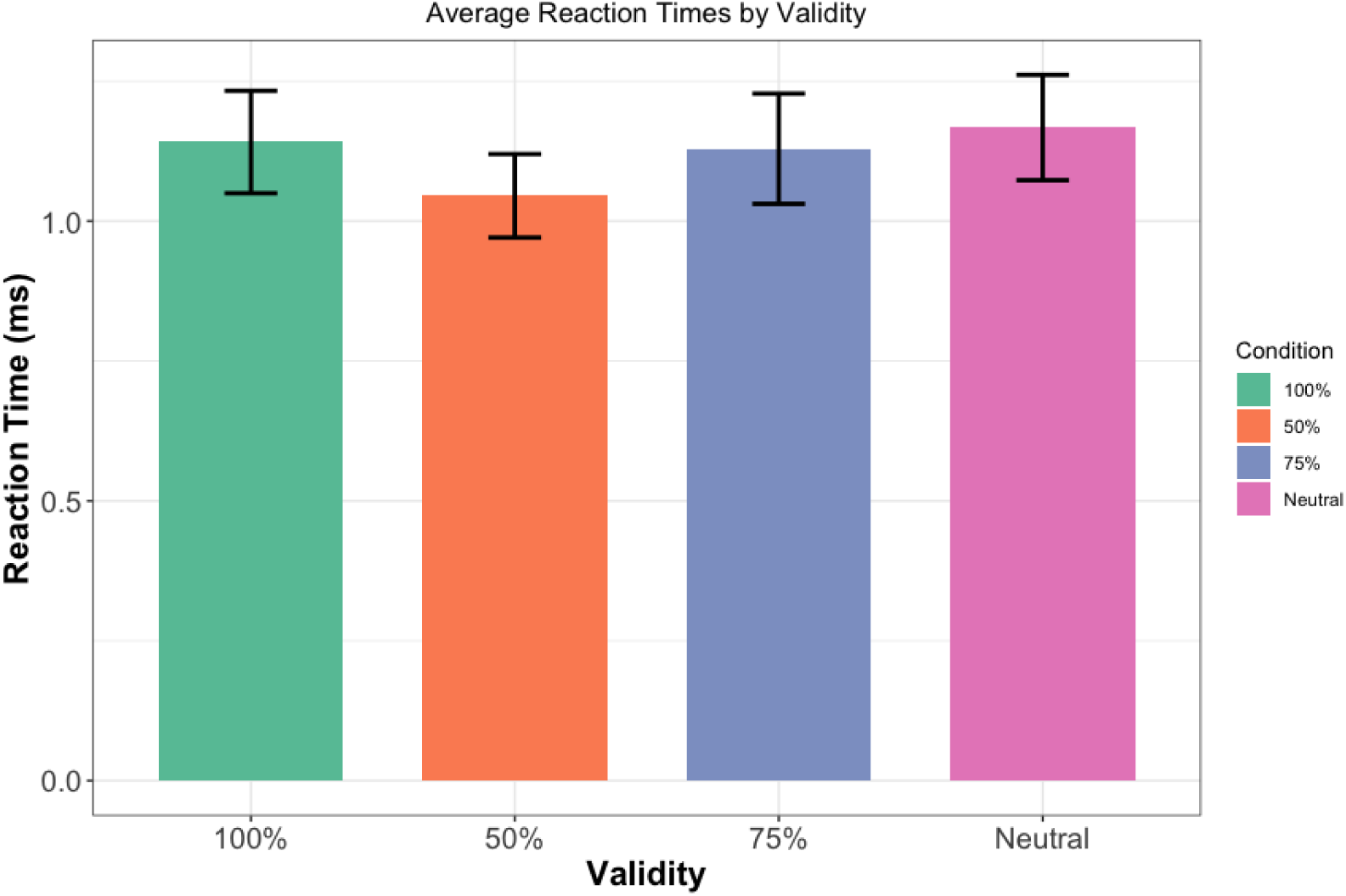
Overall reaction times for different validity conditions in emotion experiment

**Figure 7:**
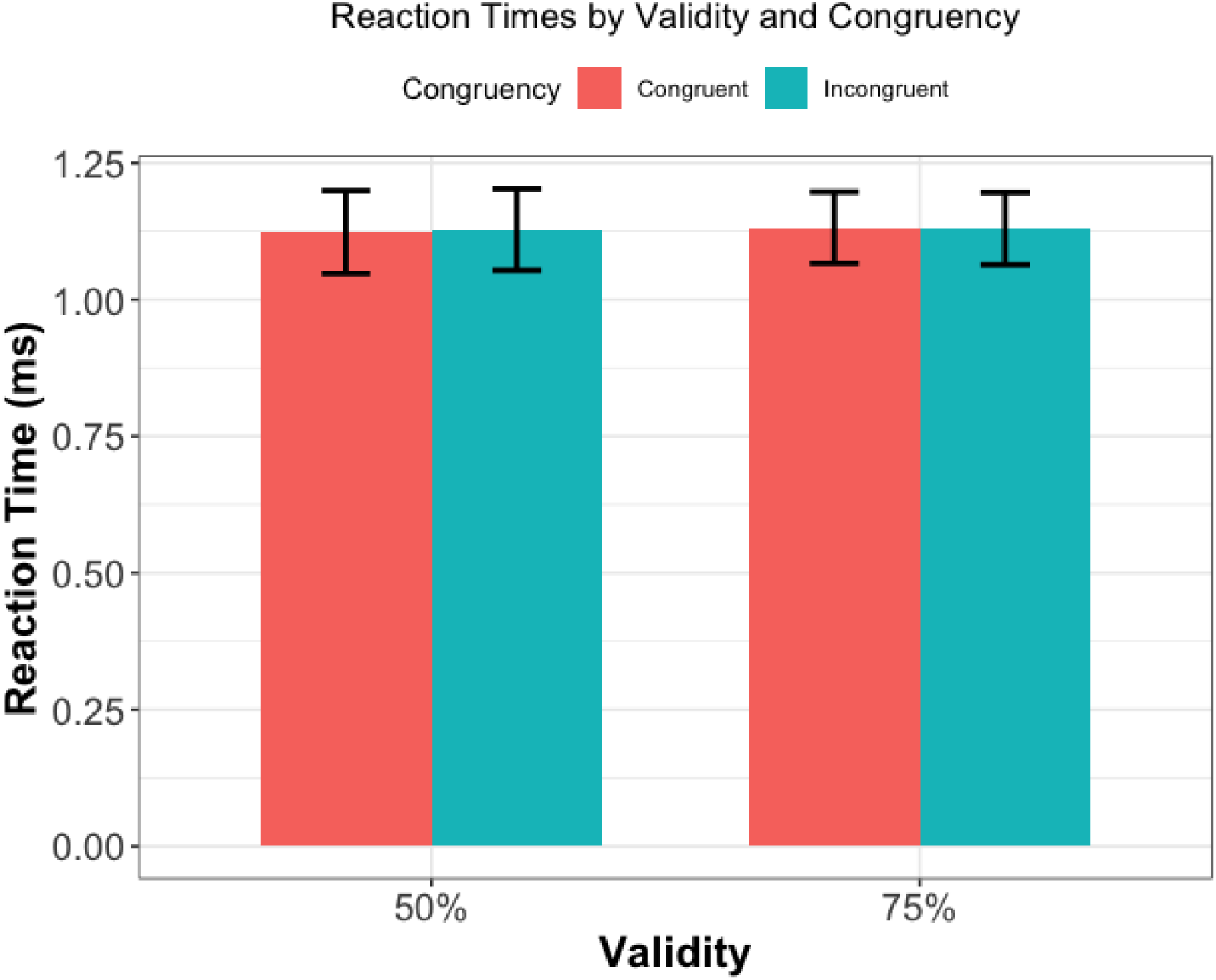
Reaction times for congruent and incongruent conditions when the validity is 50% and 75% in gender experiment

**Figure 8:**
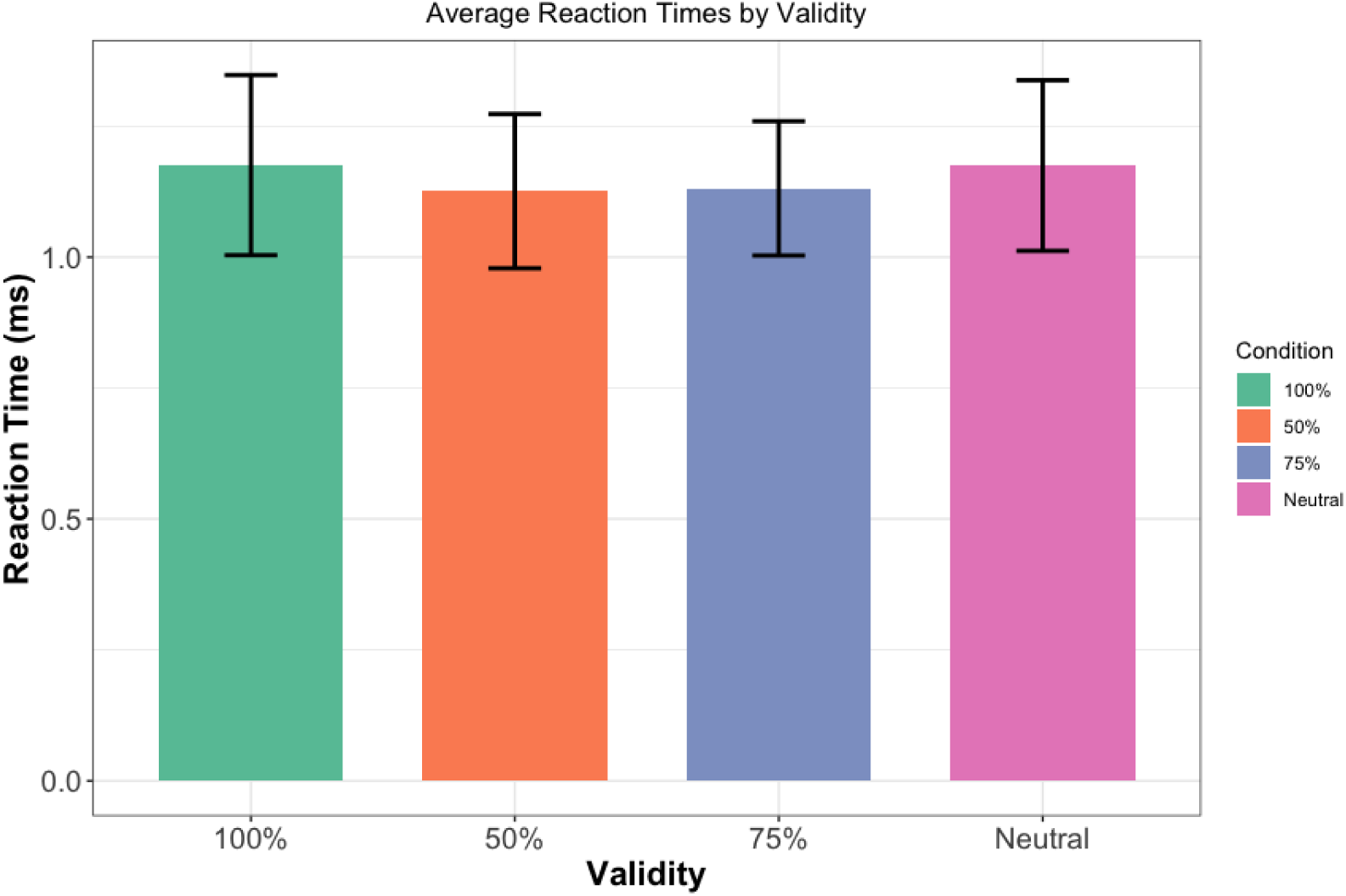
Overall reaction times for different validity conditions in gender experiment

### 3.3. Gender Experiment

In gender experiment, our 2 (Congruency) x 2 (Validity) ANOVA on reaction time data yielded no significant main effect for either congruency (F(1,20) = 0.124, p = 0.728) or validity (F(1,20) = 0.006, p = 0.939). Additionally, there was no significant interaction between congruency and validity (F(1,20) = 0.544, p = 0.470) (Figure:7).

Similar to the analysis approach of action and emotion experiments, we conducted a one-way ANOVA across validity levels for gender experiment The results showed no significant differences between the validity levels (F(3,60) = 0.473, p = 0.702)(Figure:8).

## 4. Discussion

The perception of biological motion stands as a pivotal mechanism, allowing organisms to discern and anticipate intentions from the actions of conspecifics. Despite the extensive body of research on the bottom-up processing of biological motion, the modulatory role of prior information on biological motion perception remains relatively underexplored.

In this study we examined if prior knowledge about specific features of biological motion affects its individuation, assessing the impact of the reliability of this prior information on its effects on perception, and evaluating the differential effects of various types of prior information about biological motion, specifically action, emotion, or gender.

Based on literature and our theoretical framework, we formulated the hypothesis that incongruent prior information would lead to delayed individuation of biological motion compared to congruent prior information, manifesting as smaller reaction times in congruent trials. Furthermore, we posited that the magnitude of the effect of prior information on perception would be modulated by its validity and that distinct categories of information would create differential perceptual effects.

Interestingly, our results did not indicate a general effect of cue validity in any cue type. However, a noteworthy pattern emerged in the action experiment, where we observed a significant congruent-incongruent effect, but only in the 75% validity session. This isolated finding contrasts with the absence of similar effects in the emotion and gender experiments, as well as in other cue validity levels within the action experiment.

### 4.1. Impact of Prior Information on Biological Motion Individuation

The predictive processing paradigm posits that prior information significantly influence task performance across various domains, such as detection or search tasks (Jabar and Anderson, 2017; Aitken et al., 2020; De Loof et al., 2016; Urgen and Boyaci, 2021). Nonetheless, the literature offers limited insight into if these predictive mechanisms interact with socially important and dynamic stimuli, like biological motion. The pressing question then arises: Does prior information modulate the detection and recognition of biological motion?

In the current study, our findings resonate well with results in the predictive processing literature. Specifically, during the cued individuation task—typically employed to gauge visual awareness—biological motion was more slowly individuated when the cue was incongruent compared to when the cue provided congruent and reasonably-reliable information related to the action of the upcoming biological motion stilmuli. This highlights not only the overarching influence of accurate prior information on behavioral, as evidenced in prior studies using more rudimentary stimuli but also underlines that such effects are also visible in socially significant stimuli and complex stimuli. Our results also imply that this predictive information directly modulates the processes through which biological motion ascends into visual awareness, a conclusion in harmony with prior works in the predictive processing domain. (De Loof et al., 2016; Melloni et al., 2011).

### 4.2. The effects of prior information are influenced by its validity

The framework of predictive processing relies heavily on the reliability of prior information. Within this framework, the brain is continually assessing the consistency between prior beliefs and incoming sensory data. When prior information aligns with sensory input, it’s deemed reliable, enabling the brain to confidently use it for generating predictions. Conversely, inconsistencies between prior beliefs and sensory data reduce the weighting of priors in subsequent neural computations, potentially hindering their effect on perception (Summerfield and De Lange, 2014; De Lange et al., 2018).

Recent empirical evidence supports this view. Behavioral manifestations influenced by expectations have been shown to emerge primarily when expectations are consistently valid (Urgen and Boyaci, 2021). Our results aligns with this perspective. We observed that, under 75% validity condition, participant responses were significantly influenced by the cues. However, with a cue validity at chance level (50%), their influence was not significant.

This nuanced interaction between cue reliability and prediction underscores the adaptive nature of neural processing. Rather than statically incorporating prior beliefs, the brain appears to dynamically adjust the weighting of these priors based on their past reliability. Such adaptive processing may serve to optimize neural resources and enhance behavioral outcomes in ever-changing environments.

In further examining the impact of validity on perceptual performance, our results did not reveal significant differences between different levels of validity. This lack of distinction might partly stem from the notable variability in performance of individuals across sessions, which occurred on different days. Such variability could obscure the nuanced effects of different validity levels on biological motion perception. Due to high performance variability of this task, future studies should be designed to eliminate this variablitiy, for instance by using incorparating validity levels in a single session, providing clearer insights into this relationship.

Interestingly, the fact that this interaction is observed in tasks as complex as biological motion detection suggests that reliability-based weighting of priors is a pervasive aspect of neural processing, spanning from simple perceptual tasks to complex cognitive functions. This underscores the importance of studying the interplay between reliability and prediction, especially in tasks that involve higher-order cognitive functions and social perception.

### 4.3. Different Types of Information Affect Performance Differently

Our results showed a notable distinction in how different forms of prior information influence biological motion perception. Whereas action-related cues significantly modulated participants’ reaction times, there was no analogous influence exerted by cues about emotion or gender. This discrepancy compels us to explore the potential underlying mechanisms.

At the heart of this differentiation might be the distinct neural processing associated with actions, gender, and emotions. Action recognition is intricately tied to the functioning of the superior temporal sulcus, premotor cortex, and parietal cortex, areas known to be pivotal in understanding and perceiving others’ movements (Orban, 2018). Conversely, the recognition of gender and emotions relies on other distinct regions of the brain. For instance, representation of gender of a face predominantly draws upon the fusiform face area and the extrastriate body area (Kaul et al., 2011). Anterior temporal pole also carries a role in processing gender infromation from observed actions (Orban, 2018). Emotional recognition, especially when discerning facial expressions, taps into the processing capabilities of the amygdala, insula, and orbitofrontal cortex (Phelps, 2006; Adolphs, 2002). The specialized roles and activation patterns of these regions may, therefore, result in varied behavioral outputs during tasks centered on biological motion perception. Furthermore, the ease with which action information is extracted from biological motion stands out. Human evolution has primed our neural circuits for swift and efficient action recognition, a skill vital for both survival and intricate social interactions (Blake and Shiffrar, 2007; Grossman and Blake, 2002). This historical impetus might render action cues particularly influential in shaping our perception. On the other hand, discerning emotions and gender from biological motion may involve more nuanced and layered processing. Emotion recognition, for example, requires an amalgamation of multiple cues, spanning from body postures to subtle facial expressions. Gender recognition, too, mandates the integration of various visual elements, like body shapes and the kinematics of movement (Johnson and Tassinary, 2005; Troje, 2002). The point-light displays (PLDs) in our study may not encapsulate all these subtleties, potentially rendering these categories less discernible and impacting reaction times. Moreover, the preliminary results from our norming experiments suggest potential ambiguities in participant perceptions when presented with emotion and gender cues in PLDs. The diminished distinctiveness of these visual cues could contribute to their reduced influence on reaction times.

## 5. Conclusion

Our exploration into the effects of prior information on biological motion perception, illuminates the multi-faceted nature of biological motion perception. It is evident that while our brain utilizes prior information to anticipate and interpret ensuing stimuli, the nature and reliability of these cues dictate their overall influence on biological motion perception. Specifically, action-related information emerges as a potent modulator of reaction times, possibly because of its evolutionary significance and distinctive neural processing pathway. In contrast, gender and emotion cues, despite their undeniable importance in social contexts, didn’t exert comparable influences within our experimental framework. Such findings, while consolidating some principles of predictive processing, also surface new avenues for exploration, particularly around the salience and complexity of different types of prior information in biological motion perception. As we explore these new questions, it becomes paramount to integrate diverse methodologies and stimuli, such as naturalistic videos of complex stimuli and neuroimaging methods, to comprehensively understand the mechanisms influencing our perception in the context of dynamic, socially-relevant stimuli.

## 6 Acknowledgements

The study was supported by a TUBITAK 3501 CAREER grant to Burcu A. Urgen (No: 119K654). The authors would like to thank Berfin Aydin, Cenk Günsel, Yasmine Khamkhami, and Beyza Gülzehra Ekinci for help with data collection.

## 8. CRediT authorship contribution statement

Huseyin O. Elmas: Conceptualization, Formal analysis, Investigation, Visualization, Writing – original draft, Writing – review & editing. Sena Er: Conceptualization, Investigation, Writing – original draft, Writing – review & editing. Ada D. Rezaki: Investigation. Aysesu Izgi: Investigation. Buse M. Urgen: Writing – review. Huseyin Boyaci: Writing – review. Burcu A. Urgen: Conceptualization, Funding acquisition, Resources Supervision, Writing – review & editing.

## References

1. Adolphs, R., 2002. Recognizing emotion from facial expressions: psychological and neurological mechanisms. Behavioral and cognitive neuroscience reviews 1, 21–62.

2. Aitken, F., Turner, G., Kok, P., 2020. Prior expectations of motion direction modulate early sensory processing. Journal of Neuroscience 40, 6389–6397.

3. Bang, J.W., Rahnev, D., 2017. Stimulus expectation alters decision criterion but not sensory signal in perceptual decision making. Scientific reports 7, 17072.

4. Barre, A., Armand, S., 2014. Biomechanical toolkit: Open-source framework to visualize and process biomechanical data. Computer methods and programs in biomedicine 114, 80–87.

5. Blake, R., Shiffrar, M., 2007. Perception of human motion. Annu. Rev. Psychol. 58, 47–73.

6. van Boxtel, J.J., Lu, H., 2013. A biological motion toolbox for reading, displaying, and manipulating motion capture data in research settings. Journal of vision 13, 7–7.

7. Bülthoff, I., Bülthoff, H., Sinha, P., 1998. Top-down influences on stereoscopic depth-perception. Nature neuroscience 1, 254–257.

8. Cavanagh, P., Labianca, A.T., Thornton, I.M., 2001. Attention-based visual routines: Sprites. Cognition 80, 47–60.

9. Cheadle, S., Egner, T., Wyart, V., Wu, C., Summerfield, C., 2015. Feature expectation heightens visual sensitivity during fine orientation discrimination. Journal of vision 15, 14–14.

10. De Agrò, M., Rößler, D.C., Kim, K., Shamble, P.S., 2021. Perception of biological motion by jumping spiders. PLoS biology 19, e3001172.

11. De Lange, F.P., Heilbron, M., Kok, P., 2018. How do expectations shape perception? Trends in cognitive sciences 22, 764–779.

12. De Loof, E., Van Opstal, F., Verguts, T., 2016. Predictive information speeds up visual awareness in an individuation task by modulating threshold setting, not processing efficiency. Vision research 121, 104–112.

13. Giese, M.A., Poggio, T., 2003. Neural mechanisms for the recognition of biological movements. Nature Reviews Neuroscience 4, 179–192.

14. Gilbert, C.D., Li, W., 2013. Top-down influences on visual processing. Nature Reviews Neuroscience 14, 350–363.

15. Grossman, E.D., Blake, R., 2002. Brain areas active during visual perception of biological motion. Neuron 35, 1167–1175.

16. Hirai, M., Senju, A., Fukushima, H., Hiraki, K., 2005. Active processing of biological motion perception: an erp study. Cognitive Brain Research 23, 387–396.

17. Hodgins, J., 2015. Cmu graphics lab motion capture database.

18. Hunt, A.R., Halper, F., 2008. Disorganizing biological motion. Journal of Vision 8, 12–12.

19. Jabar, S.B., Anderson, B., 2017. Not all probabilities are equivalent: Evidence from orientation versus spatial probability learning. Journal of experimental psychology: human perception and performance 43, 853.

20. Johansson, G., 1973. Visual perception of biological motion and a model for its analysis. Perception & psychophysics 14, 201–211.

21. Johnson, K.L., Tassinary, L.G., 2005. Perceiving sex directly and indirectly: Meaning in motion and morphology. Psychological Science 16, 890–897.

22. Kaul, C., Rees, G., Ishai, A., 2011. The gender of face stimuli is represented in multiple regions in the human brain. Frontiers in Human Neuroscience 4, 238.

23. Kok, P., Jehee, J.F., De Lange, F.P., 2012. Less is more: expectation sharpens representations in the primary visual cortex. Neuron 75, 265–270.

24. Kok, P., Mostert, P., De Lange, F.P., 2017. Prior expectations induce prestimulus sensory templates. Proceedings of the National Academy of Sciences 114, 10473–10478.

25. Lange, J., Georg, K., Lappe, M., 2006. Visual perception of biological motion by form: A template-matching analysis. Journal of vision 6, 6–6.

26. Ma, Y., Paterson, H.M., Pollick, F.E., 2006. A motion capture library for the study of identity, gender, and emotion perception from biological motion. Behavior research methods 38, 134–141.

27. Mather, G., Radford, K., West, S., 1992. Low-level visual processing of biological motion. Proceedings of the Royal Society of London. Series B: Biological Sciences 249, 149–155.

28. Melloni, L., Schwiedrzik, C.M., Müller, N., Rodriguez, E., Singer, W., 2011. Expectations change the signatures and timing of electrophysiological correlates of perceptual awareness. Journal of Neuroscience 31, 1386–1396.

29. Omori, E., Watanabe, S., 1996. Discrimination of johansson’s stimuli in pigeons. Int J Comp Psychol 9, 92.

30. Orban, G.A., 2018. Action observation as a visual process: Different classes of actions engage distinct regions of human ppc, in: CULTURAL PATTERNS AND NEUROCOGNITIVE CIRCUITS II: East–West Connections. World Scientific, pp. 1–32.

31. Parasuraman, R., de Visser, E., Clarke, E., McGarry, W.R., Hussey, E., Shaw, T., Thompson, J.C., 2009. Detecting threat-related intentional actions of others: effects of image quality, response mode, and target cuing on vigilance. Journal of experimental psychology: applied 15, 275.

32. Peelen, M.V., Berlot, E., de Lange, F.P., 2023. Predictive processing of scenes and objects. Nature Reviews Psychology URL: https://www.nature.com/articles/s44159-023-00254-0, doi:10.1038/s44159-023-00254-0.

33. Phelps, E.A., 2006. Emotion and cognition: insights from studies of the human amygdala. Annu. Rev. Psychol. 57, 27–53.

34. Qualtrics, 2023. Qualtrics. https://www.qualtrics.com. URL: https://www.qualtrics.com. version used: Sep 2021. Available at: https://www.qualtrics.com.

35. Saygin, A.P., Chaminade, T., Ishiguro, H., Driver, J., Frith, C., 2012. The thing that should not be: predictive coding and the uncanny valley in perceiving human and humanoid robot actions. Social cognitive and affective neuroscience 7, 413–422.

36. Simion, F., Regolin, L., Bulf, H., 2008. A predisposition for biological motion in the newborn baby. Proceedings of the National Academy of Sciences 105, 809–813.

37. Spaak, E., Peelen, M.V., de Lange, F.P., 2022. Scene context impairs perception of semantically congruent objects. Psychological Science 33, 299–313.

38. Summerfield, C., De Lange, F.P., 2014. Expectation in perceptual decision making: neural and computational mechanisms. Nature Reviews Neuroscience 15, 745–756.

39. Thornton, I.M., Pinto, J., Shiffrar, M., 1998. The visual perception of human locomotion. Cognitive Neuropsychology 15, 535–552.

40. Thornton, I.M., Rensink, R.A., Shiffrar, M., 2002. Active versus passive processing of biological motion. Perception 31, 837–853.

41. Troje, N.F., 2002. Decomposing biological motion: A framework for analysis and synthesis of human gait patterns. Journal of vision 2, 2–2.

42. Troje, N.F., 2008. Retrieving information from human movement patterns. Understanding events: How humans see, represent, and act on events 1, 308–334.

43. Troje, N.F., Basbaum, A., 2008. Biological motion perception. The senses: A comprehensive reference 2, 231–238.

44. Urgen, B.A., Kutas, M., Saygin, A.P., 2018. Uncanny valley as a window into predictive processing in the social brain. Neuropsychologia 114, 181–185.

45. Urgen, B.A., Miller, L.E., 2015. Towards an empirically grounded predictive coding account of action understanding. Journal of Neuroscience 35, 4789– 4791.

46. Urgen, B.A., Saygin, A.P., 2020. Predictive processing account of action perception: Evidence from effective connectivity in the action observation network. Cortex 128, 132–142.

47. Urgen, B.M., Boyaci, H., 2021. Unmet expectations delay sensory processes. Vision Research 181, 1–9.

48. Vallortigara, G., Regolin, L., Marconato, F., 2005. Visually inexperienced chicks exhibit spontaneous preference for biological motion patterns. PLoS biology 3, e208.

49. Watson, A.B., Pelli, D.G., 1983. Quest: A bayesian adaptive psychometric method. Perception & psychophysics 33, 113–120.

